# ImmuneLens: linking transcriptional states and TCR clonotypes through disentangled multimodal learning

**DOI:** 10.64898/2026.09.08.749998

**Authors:** Zhijian Duan, Yi Wang, Cuiping Li, Guochao Li, Yongrong Cao, Xue Bai, Fei Yang, Shuhui Song

## Abstract

Single-cell multi-omics technologies simultaneously capture the transcriptome and TCR sequence of T cells, providing an opportunity to study the relationship between transcriptional states and clonal architectures. However, jointly modeling the relationships between transcriptional states and TCR sequences while preserving modality-specific information remains challenging. Here, we present ImmuneLens, an interpretable multimodal representation learning framework designed for paired single-cell transcriptome and TCR sequence data. ImmuneLens supports the construction of a transferable multi-cohort immune reference atlas and enables unsupervised mapping of external query data. The complementarity between GEX and TCR information improves the stability of antigen-specificity prediction. In neoadjuvant immunotherapy cohorts, ImmuneLens resolves response-associated T cell heterogeneity and reveals links between clonal expansion and CD8 T cell functional states. Overall, ImmuneLens provides a transferable and interpretable framework for linking transcriptional states, TCR clonal architecture, antigen recognition, and disease phenotypes in single-cell immunology.

## INTRODUCTION

T cells are central effector units of the adaptive immune system and play key roles in infection clearance, tumor immune surveillance, and the maintenance of autoimmune homeostasis ^1^. The functional states of T cells are determined not only by their transcriptomes but also by antigen-specific signals mediated by the T cell receptor (TCR) ^2^. With the development of single-cell sequencing technologies, the simultaneous acquisition of transcriptomic and TCR sequence information at the same single-cell level has provided unprecedented opportunities to systematically dissect the relationships among T cell phenotypes, functional states, and antigen recognition.

In recent years, large-scale single-cell RNA sequencing has been widely applied to immune cell atlas construction and cell state characterization^3–5^, whereas TCR sequencing has provided key clues for tracking clonal expansion, antigen-driven selection, and immune response dynamics^6^. Joint analysis of these two modalities is expected to connect the molecular states of T cells with their immunological functions at the clonal level^7^, thereby deepening our understanding of how immune responses are organized in complex disease contexts, including infection, autoimmune diseases, and tumors.

Recently, deep learning methods have become increasingly important tools for single-cell data analysis because of their strong nonlinear modeling capacity^8^. A variety of deep learning models have been developed for transcriptomic analysis. For example, models such as scCapsNet^9^ and scBERT^10^ can learn cell state representations from high-dimensional and sparse gene expression data. In the field of TCR analysis, methods such as DeepTCR^11^ and TCR-BERT^12^ use sequence models to learn sequence representations relevant to antigen specificity. However, these methods are generally limited to unimodal modeling and do not explore the intrinsic associations between transcriptomic states and TCR sequences.

Recent methods have approached the joint analysis of single-cell transcriptomic and TCR sequence data from different perspectives. CoNGA compares neighborhood structures across transcriptomic and TCR spaces^13^, Tessa links TCR sequence similarity with transcriptional states^14^, and deep generative models such as scNAT^15^, mvTCR^16^, and MIST^17^ learn latent representations from paired GEX and TCR data. These approaches have substantially advanced multimodal T cell analysis. However, fine-grained interactions between gene expression features and individual TCR sequence positions, together with the explicit separation of shared and modality-specific information, remain less extensively explored.

In this study, we present ImmuneLens, a deep learning framework for paired single-cell transcriptomic and TCR sequence data. ImmuneLens explicitly models GEX-private, TCR-private, and cross-modal shared representations. A multilayer cross-attention mechanism captures fine-grained interactions between gene expression and TCR sequence features, while gated fusion regulates the contribution of each modality to the shared representation.

Based on this multimodal representation learning framework, we systematically evaluate ImmuneLens from three aspects: multi-cohort immune atlas construction, antigen-specificity recognition, and tumor immunotherapy-associated state characterization. First, we evaluate the GEX branch of ImmuneLens for constructing a multi-cohort immune reference atlas and performing unsupervised query mapping of external data. Next, we assess the complementary roles of GEX and TCR information through an antigen recognition prediction task. Finally, we apply ImmuneLens to tumor immunotherapy data to examine the utility of the fused representation in resolving clinically relevant T cell states. The results show that ImmuneLens provides transferable GEX representations for cross-dataset integration and query mapping, together with interpretable multimodal representations for antigen-associated signal capture and disease-associated T cell state characterization.

In summary, ImmuneLens provides a unified multimodal modeling framework for systematically dissecting the complex relationships among T cell transcriptional states, TCR clonal architecture, and antigen recognition at single-cell resolution, thereby laying a methodological foundation for immunological research and immunotherapy-related applications.

## RESULTS

### Overview of the ImmuneLens multimodal framework

ImmuneLens is a multimodal representation learning framework for the joint modeling of single-cell transcriptomic and TCR sequence information (**Figure 1a**), with the detailed network architecture and cross-attention module provided in **Figure S1**. This framework separately encodes gene expression profiles and TCR CDR3 sequences, and learns the interactions between GEX and TCR sequences through a cross-modal cross-attention mechanism. To prevent the fused representation from being dominated by a single modality, ImmuneLens further introduces shared and private representation disentanglement in the latent space, encoding immune states jointly supported by GEX and TCR into a cross-modal shared space while retaining modality-specific information in private spaces. Through this design, ImmuneLens can preserve modality-specific information such as transcriptional states, TCR sequences, and clonal architecture, while also learning a fused representation that integrates GEX and TCR information.

**Figure 1.**
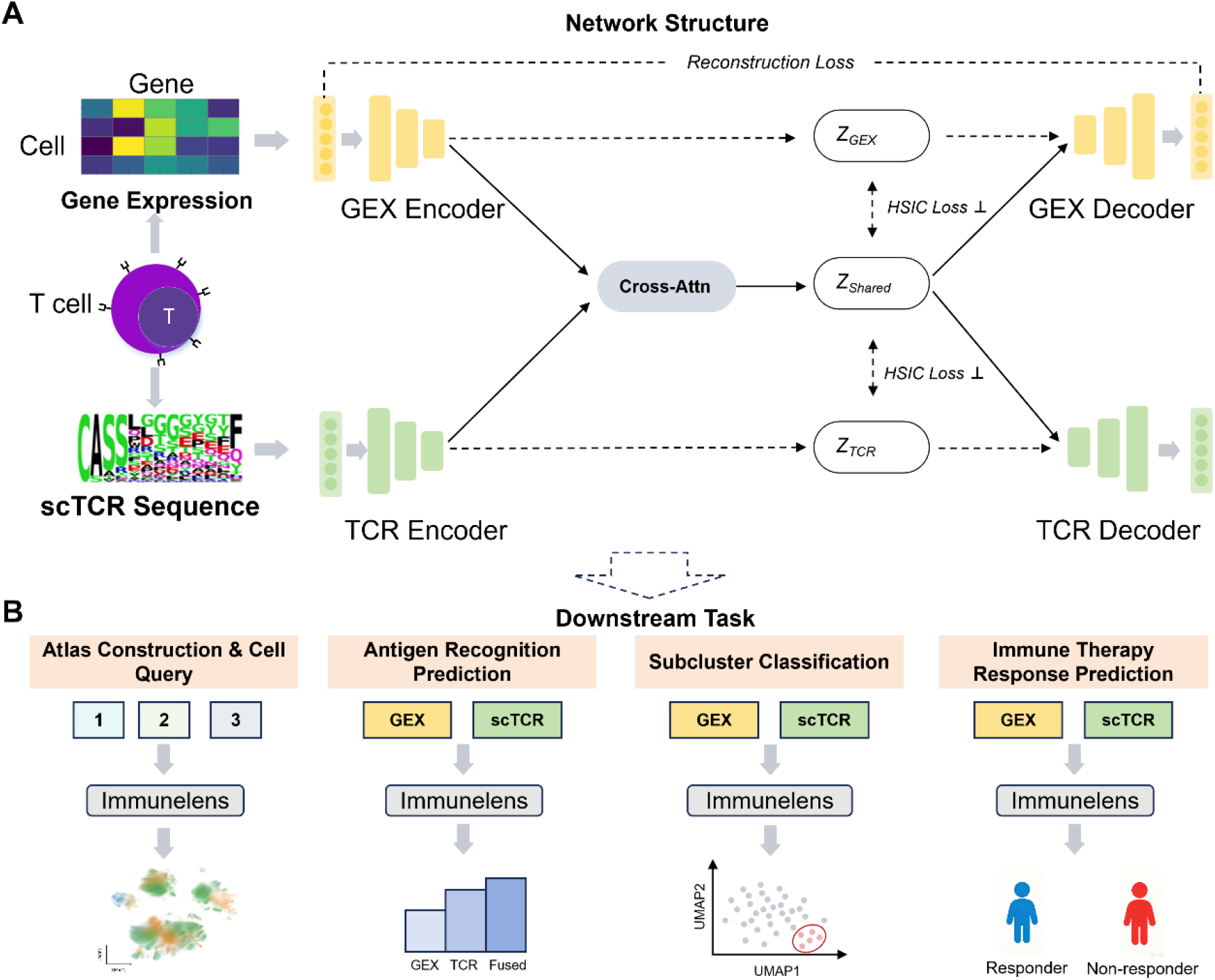
Overview of the ImmuneLens model framework and downstream tasks. **(A)** The multimodal disentangled representation learning framework of ImmuneLens for paired single-cell transcriptome and TCR sequence data. **(B)** Schematic illustration of downstream analysis tasks based on ImmuneLens representations. See also Figure S1.

Based on this disentangled multimodal representation, ImmuneLens can support immune analysis tasks at different scales (**Figure 1b**). First, we constructed a multi-cohort pan-immune reference atlas and performed unsupervised query mapping on an external skin cancer dataset. Second, we examined the complementarity of GEX and TCR information in identifying T cell antigen responses through an antigen-specificity recognition task. Third, we applied ImmuneLens to a lung adenocarcinoma neoadjuvant immunotherapy cohort and further resolved subclusters in the fused representation space that were differentially associated with pathological responders and non-responders within GEX-only-defined tumor-reactive/exhaustion-like T cell states. Finally, we constructed an immunotherapy response prediction task in lung squamous cell carcinoma and analyzed highlighted clones identified from the fused representation, demonstrating that ImmuneLens can capture treatment response-associated CD8 T cell states with clonal expansion backgrounds and functional features.

### ImmuneLens constructs a multi-cohort immune reference atlas and supports external data projection

Immune cell states are shaped by both conserved biological programs and dataset-specific batch effects. Therefore, a unified reference space is required for cross-dataset comparison of immune cell states and cell type annotation. Here, we evaluated whether the GEX encoder of ImmuneLens could construct a unified reference space across cohorts using single-cell transcriptomic data from colorectal cancer (CRC) ^18^, non-small cell lung cancer (NSCLC)^19^, and COVID-19^20^. The major immune cell types shared across the three disease cohorts included B lineage cells, CD4 T cells, CD8 T cells, DCs, macrophages, monocytes, NK/ILC cells, plasma cells, and γδ T cells, comprising approximately 3.35 million immune cells. Although the proportions of individual immune cell types varied across cohorts, the major immune cell lineages were broadly represented in the combined reference dataset (**Figure S2a**).

We compared the cross-datasets integration performance by three different embedding spaces: Raw, scVI^21^, and ImmuneLens (**Figure 2a**). In the raw expression space (Raw), cell distributions were clearly influenced by dataset origin, with significant separation between different cohorts, and batch effects had a strong influence on the overall structure. The mainstream integration method scVI could partially alleviate this effect, but residual cohort-associated separation structures could still be observed. In contrast, ImmuneLens achieved more consistent cross-cohort cell mixing while preserving the spatial organization of major immune cell types. Quantitative evaluation (**Figure 2b**) further supported the above results, showing that ImmuneLens outperformed Raw and scVI in overall integration performance while still preserving the consistency of cell type structure.

**Figure 2.**
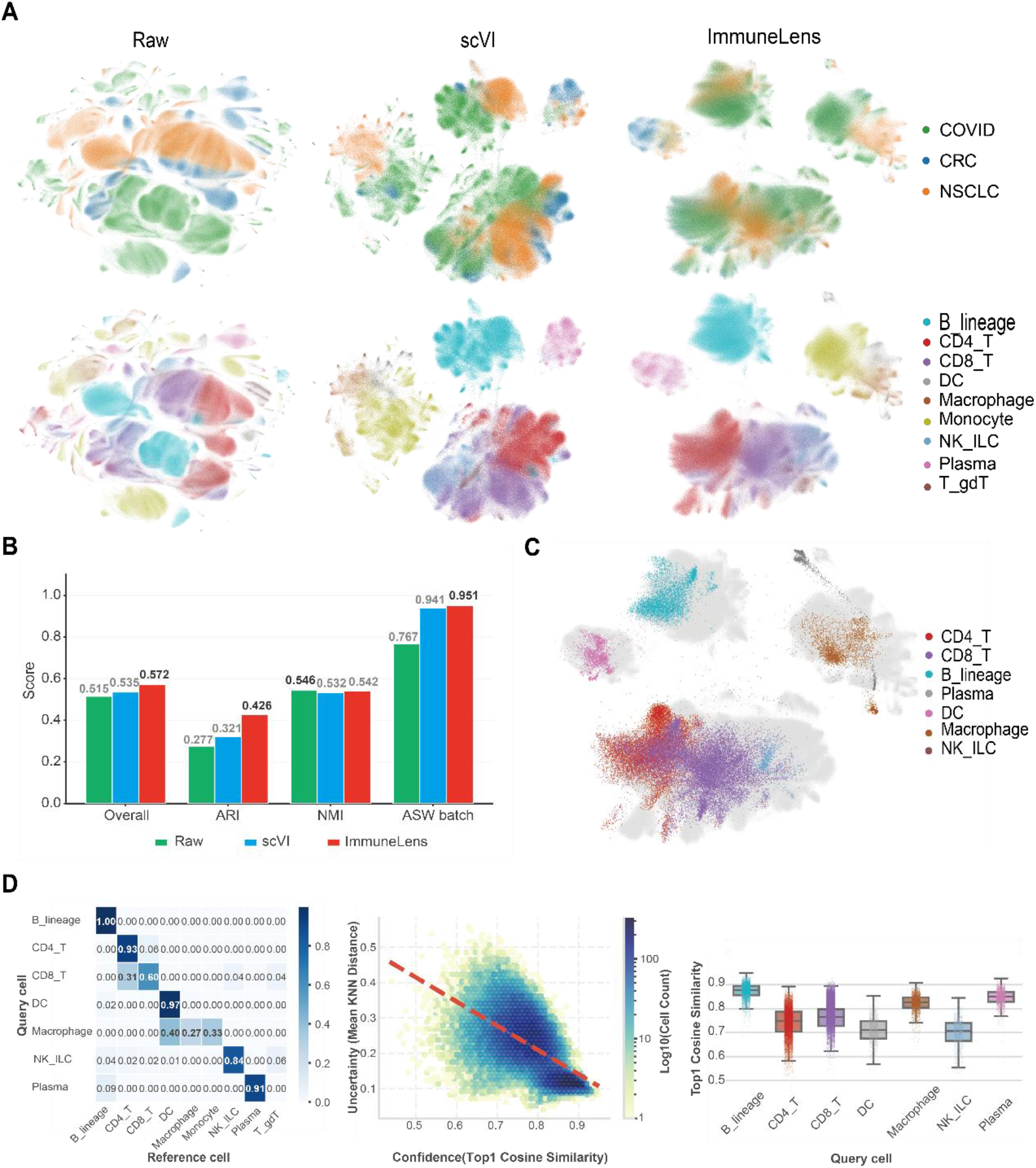
Construction of a multi-cohort immune reference atlas and query mapping of external data. **(A)** UMAP visualization of the Raw, scVI, and ImmuneLens embedding spaces. The upper row is colored by dataset origin, and the lower row is colored by major cell type. **(B)** Integration performance of different methods, evaluated using Overall, ARI, NMI, and ASW batch metrics. Higher values indicate better integration performance. **(C)** Projection of BCC cells onto the ImmuneLens reference atlas. Gray indicates reference cells, and colored points indicate BCC query cells. **(D)** Evaluation of BCC query mapping by annotation consistency, confidence–uncertainty relationship, and cell type-specific confidence distributions. See also Figure S2.

To evaluate the query mapping ability of this reference atlas for external tumor immune cells, we further introduced a basal cell carcinoma (BCC)^22^ dataset as an independent query dataset. The BCC dataset contained seven major immune cell types with clearly distinguishable transcriptional structures in the original embedding space (**Figure S2b**).

We performed query mapping of external BCC data using the pretrained ImmuneLens GEX encoder. BCC cells were stably projected onto their corresponding immune cell regions in the ImmuneLens GEX reference atlas (**Figure 2c**). Quantitative evaluation showed high mapping consistency between BCC query cells and reference cell types (**Figure 2d**). Prediction confidence was moderately negatively correlated with mapping uncertainty (Spearman’s ρ = −0.608, P < 0.001). Median prediction confidence varied across cell types, ranging from 0.706 in NK/ILC cells to 0.877 in B-lineage cells.

Overall, these results indicate that the multi-cohort immune reference atlas constructed using the ImmuneLens GEX encoder can not only effectively integrate multi-cohort data but also serve as a stable reference space to support unsupervised query mapping, projection, and consistency evaluation of external tumor immune cells. This framework provides a basis for constructing immune reference atlases across multiple datasets, as well as for the rapid localization and annotation of new samples within an existing immune space.

### ImmuneLens improves antigen recognition prediction through complementary GEX and TCR information

T cell recognition of antigens is not solely determined by TCR sequences but is also influenced by the cellular state. However, the cooperative roles of transcriptional programs and TCR sequences in antigen recognition remain to be fully elucidated. Here, we used a publicly available 10x Genomics multimodal CD8 T cell dataset from four healthy donors to evaluate whether ImmuneLens can leverage complementary information from single-cell transcriptional states and TCR sequences for antigen recognition prediction. This dataset provides paired single-cell gene expression profiles, TCR V(D)J sequences, and antigen-binding labels derived from dCODE Dextramer assays.

Based on the binarized antigen-binding matrix, we constructed per-cell labels and defined the prediction task as a binary classification over cell–antigen pairs. Using TCR-BERT^12^ and ESM2^23^ as TCR sequence encoders, we compared the ImmuneLens fused model against GEX-only and TCR-only unimodal baselines across four donor datasets. The results showed that, regardless of whether TCR-BERT or ESM2 was used as the TCR encoder, the fused model achieved the highest AUPRC in most donors and showed more stable predictive performance overall (**Figure 3a**). The relative performance patterns were consistent across multiple random seeds, with the fused models generally maintaining competitive or superior performance across donors (**Figure S3**). Compared with the best unimodal model, the fused model improved performance in Donor1, Donor2, and Donor4, demonstrating that transcriptional states and TCR sequences are not completely redundant but instead provide complementary information for antigen recognition. Notably, in Donor3, GEX-only performed slightly better than the fused model, suggesting that the dominant modality of antigen-specific signals may differ across different individuals. To further dissect the relative contributions of GEX and TCR information, we performed modality-shuffling experiments, that is one modality was randomly shuffled while the other was kept unchanged to observe changes in model performance. The results showed that there was no consistent pattern among donors in their use of the two modalities. Donor2 and Donor3 illustrate this difference. The AUPRC remained relatively high after GEX shuffling for Donor2, implying that TCR signals alone were highly discriminative. For Donor3, however, TCR shuffling caused only a marginal performance drop, suggesting that GEX itself carried strong antigen-related information(**Figure 3b**). Further class imbalance analysis revealed that raising the negative-to-positive sample ratio led to AUPRC reductions across all tested models. However, our fused model consistently outperformed the others, showing the slowest degradation and sustaining superior performance even under severely imbalanced conditions (**Figure 3c**).

**Figure 3.**
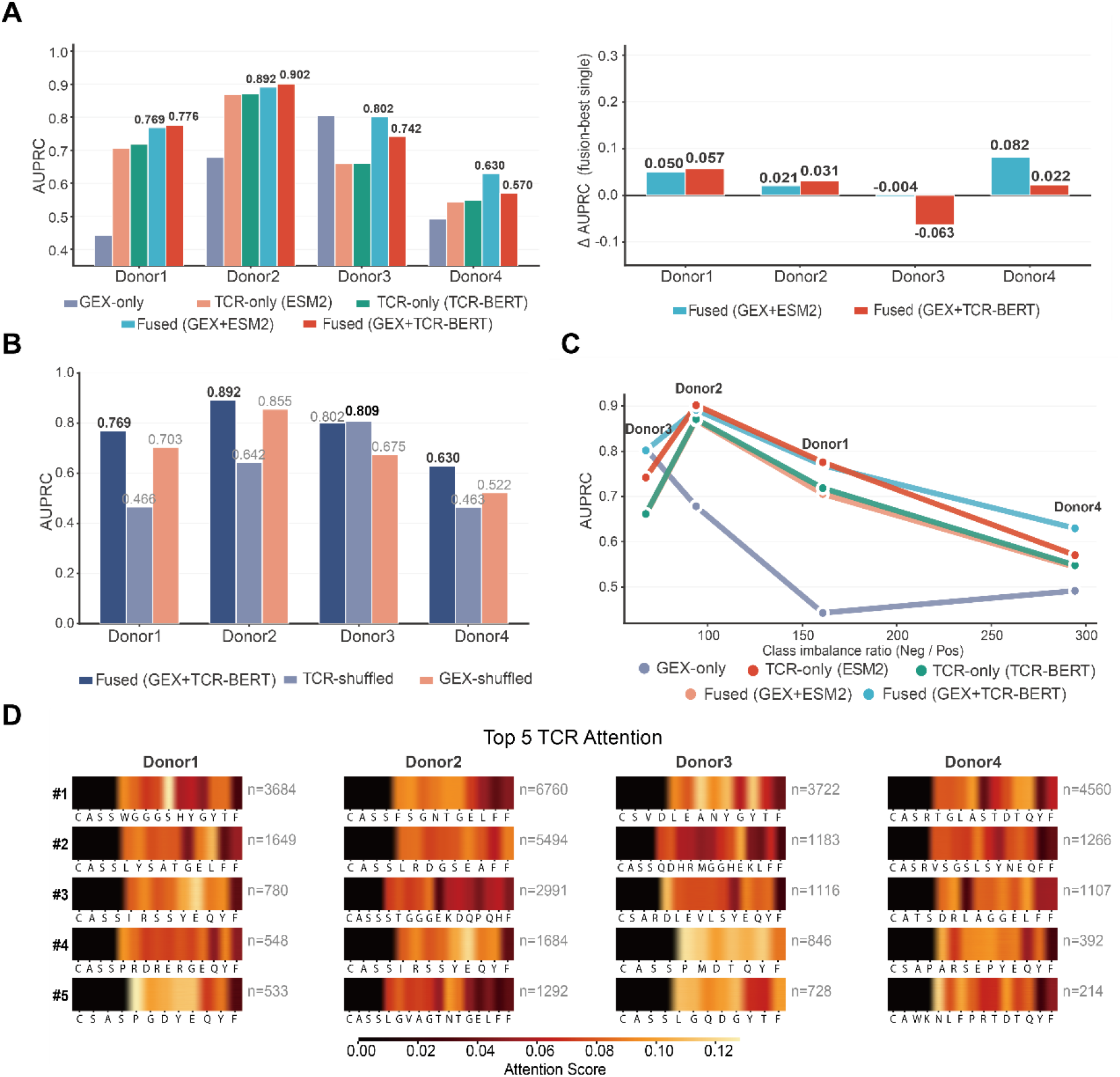
ImmuneLens integrates GEX and TCR information for antigen-specific T cell recognition. **(A)** Comparison of antigen recognition prediction performance between the fused model and the GEX-only and TCR-only unimodal baseline models across four donor datasets using TCR-BERT and ESM2 as TCR encoders. The left panel shows test AUPRC, and the right panel shows the AUPRC gain of the fused model relative to the best unimodal model. **(B)** Modality-shuffling analysis. One modality was randomly shuffled while the other was kept unchanged to evaluate the model dependence on GEX or TCR signals. **(C)** Changes in test AUPRC of each model under different degrees of class imbalance. Each point represents one donor, and lines indicate the performance changes of the same model under different negative-to-positive sample ratios. **(D)** Attention visualization of representative TRB clonotypes across different donors. CDR3 attention heatmaps are shown for the top five clonotypes in each donor. Colors indicate attention scores, and n indicates the number of cells contained in the corresponding clonotype. See also Figure S3.

We further evaluated the interpretability of the fused model by analyzing the attention distribution of TRB sequences. Visualization of top-ranked clonotypes from each donor showed that our fused model did not distribute attention uniformly across the entire CDR3 sequence, but instead focused on specific amino acid positions, forming relatively clear high-attention regions (**Figure 3d**). These high-attention regions were mainly located in the middle of the CDR3 sequence and were more likely to correspond to key sequence segments involved in antigen recognition^24^. In addition, cells within the same clonotype exhibited similar attention patterns, indicating that ImmuneLens learned stable clonotype-associated sequence features rather than incidental memorization of individual cells.

These results suggest that antigen recognition is not determined solely by TCR sequences, but may also be influenced by the transcriptional states of T cells. By jointly modeling GEX and TCR sequence information, ImmuneLens can more stably capture antigen recognition-associated signals across different donors and class imbalance conditions, while providing a degree of sequence-level interpretability.

### ImmuneLens resolves response-associated T cell subclusters by integrating transcriptional and TCR information

Transcriptome-based analyses alone may not fully capture the functional heterogeneity within a given cell population. For example, T cell populations with similar transcriptomic profiles may exhibit distinct clinical response patterns during immunotherapy. We therefore investigated whether the joint analysis of single-cell gene expression and T cell receptor sequences could further resolve T cell heterogeneity associated with immunotherapy response. We analyzed a neoadjuvant anti-PD-1 cohort of 60 patients with lung adenocarcinoma (LUAD)^19^, for whom paired single-cell RNA sequencing (GEX) and single-cell TCR sequencing data were available (**Figure 4a**). Patients were classified as responders or non-responders based on pathological response. We then generated transcriptome-based GEX representations and joint GEX–TCR representations using ImmuneLens, and compared transcriptome-only clustering with fused clustering in their ability to resolve treatment-associated cellular states.

**Figure 4.**
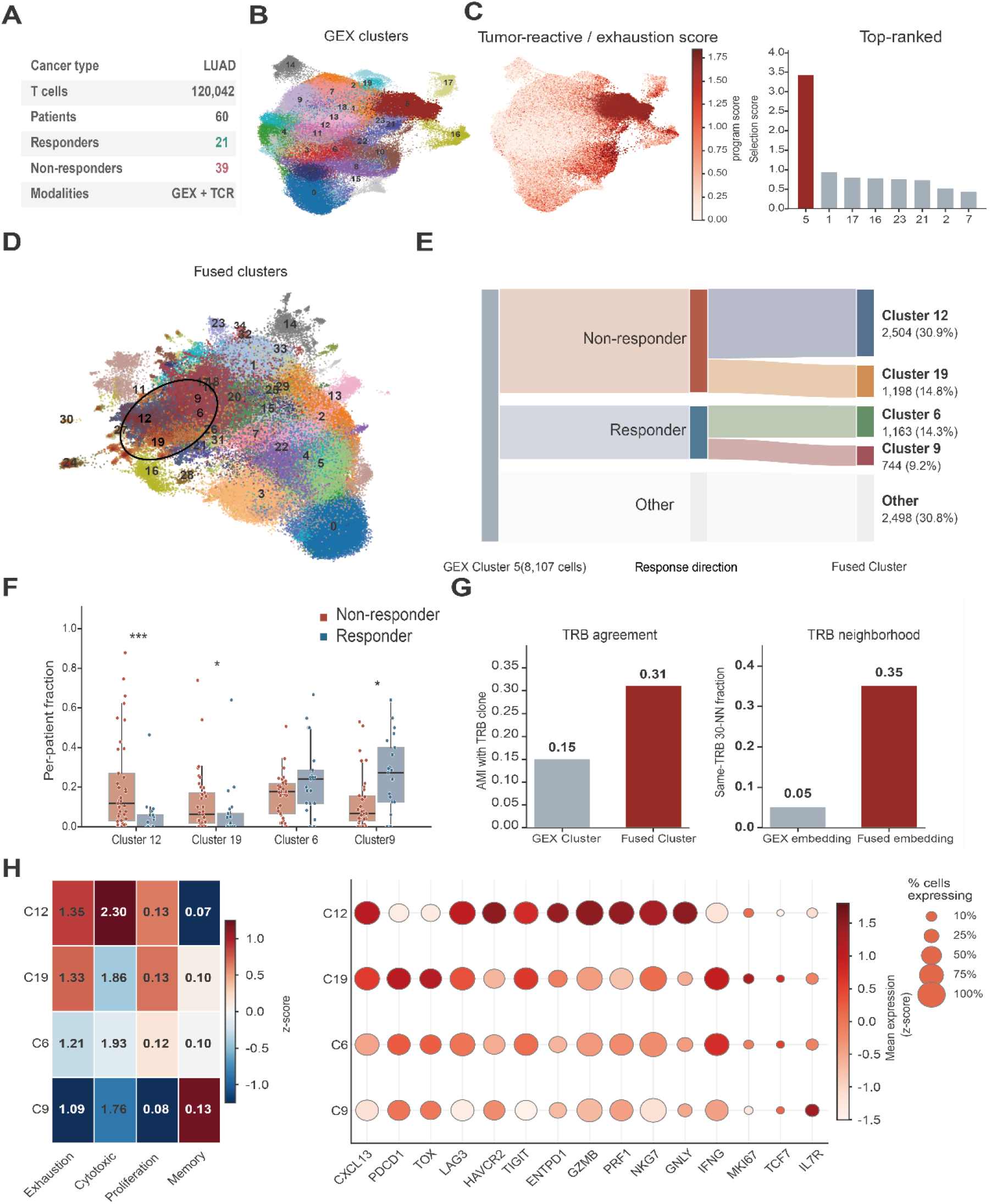
ImmuneLens resolves pathological response heterogeneity within GEX-defined T cell states. **(A)** Overview of the LUAD training dataset. **(B)** Leiden clustering results in the GEX representation space. **(C)** Distribution of the tumor-reactive/exhaustion program score on the GEX UMAP and ranking of selection scores across GEX clusters. GEX cluster 5 was selected for subsequent analysis. **(D)** Leiden clustering results in the ImmuneLens fused representation space. **(E)** Distribution of GEX cluster 5 cells across fused subclusters and their treatment response directions. **(F)** Comparison of the composition proportions of the top four fused subclusters by cell number between responder and non-responder samples. **(G)** Comparison of TRB clonal architecture between GEX-only and fused representations, including TRB clonotype agreement and TRB neighborhood. **(H)** Functional program scores and marker gene expression of representative fused subclusters. See also Figure S4.

Leiden clustering of the GEX representations identified 23 T cell clusters in the LUAD neoadjuvant anti-PD-1 cohort **(Figure 4b**). We then scored each cluster for a predefined Tex/tumor-reactive transcriptional program associated with response to immune checkpoint blockade^25–27^. GEX cluster 5 had the highest program score and was selected for subsequent ImmuneLens analysis (**Figure 4c; Figure S4a,b**). By incorporating TCR-derived information into the GEX representation, ImmuneLens further partitioned the GEX-defined cluster 5 into multiple fused subclusters that were not resolved in the transcriptome-only space **(Figure 4d**). The four largest fused clusters—FCluster 12, FCluster 19, FCluster 6, and FCluster 9—together accounted for approximately 69% of the cells in GEX cluster 5 and were retained for further analysis. These fused subclusters showed distinct response-associated abundance patterns. FCluster 12 and FCluster 19 were associated with non-response, whereas FCluster 6 and FCluster 9 were associated with response (**Figure 4e**). At the patient level, FCluster 12 and FCluster 19 were significantly more abundant in non-responders, whereas FCluster 9 was significantly more abundant in responders. FCluster 6 showed a responder-associated trend (**Figure 4f**). Thus, the GEX-defined tumor-reactive/exhaustion-like population comprised fused subclusters with different associations with pathological response. Fused cluster assignments also showed greater agreement with TRB clonotypes than GEX-only cluster assignments. In addition, cells in the fused embedding had a higher proportion of neighboring cells from the same TRB clonotype, and this pattern was consistent across different neighborhood sizes **(Figure 4g; Figure S4e,f**). The fused representation was therefore more closely aligned with TRB clonal structure than the GEX-only representation^28^.

We further compared the functional programs and marker gene expression of FCluster 12, FCluster 19, FCluster 6, and FCluster 9 (**Figure 4h**). All four fused clusters showed tumor-reactive/exhaustion-associated features but differed in their exhaustion, cytotoxicity, proliferation, and memory programs. FCluster 12 and FCluster 19 had higher exhaustion and cytotoxicity scores, together with higher expression of *PDCD1, TOX, LAG3, HAVCR2, TIGIT, ENTPD1*, and cytotoxicity-associated genes. FCluster 9 had a higher memory score and higher *IL7R* expression, whereas FCluster 6 showed an intermediate functional profile. The four fused clusters therefore represented functionally distinct states within the GEX-defined tumor-reactive/exhaustion-like population.

### ImmuneLens predicts immunotherapy response and identifies response-associated CD8 T cell clones

Although T cell clonal expansion has been associated with immunotherapy response, whether TCR-derived information provides predictive value beyond transcriptomic profiling at the single-cell level remains unclear. We next analyzed the LUSC subset of the same broader NSCLC study cohort, comprising 162 patients and 151,935 CD8 T cells with paired single-cell GEX and TCR sequencing data (**Figure 5a**). Patients were classified as responders or non-responders using the same pathological response criteria as in the LUAD analysis. We then compared ImmuneLens with GEX-only and TCR-only models for sample-level response classification.

**Figure 5.**
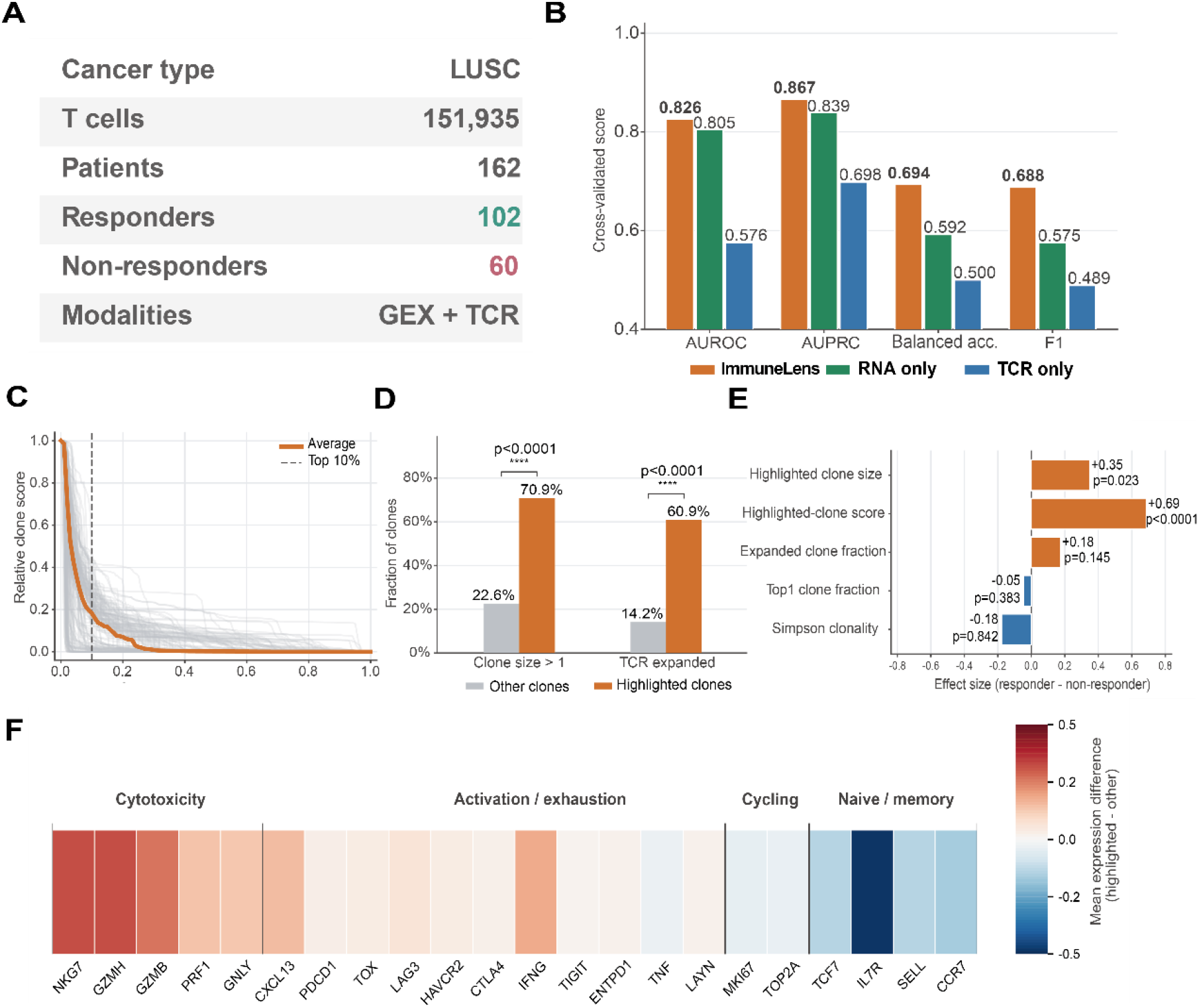
ImmuneLens predicts pathological response to neoadjuvant immunotherapy and resolves highlighted clonal CD8 T cell states. **(A)** Overview of the LUSC training dataset. **(B)** Predictive performance of the GEX-only, TCR-only, and ImmuneLens models in the immunotherapy classification task. **(C)** Relative score distribution of clones within each sample after ranking from high to low according to model scores. Gray lines indicate individual samples, the orange line indicates the median sample, and the dashed line indicates the top 10% threshold within each sample. The top 10% of clones were defined as highlighted clones. **(D)** Comparison of clonal expansion proportions between highlighted clones and other clones. Clonal expansion was evaluated using two definitions: clone size > 1 and the TCR expanded label. **(E)** Comparison of highlighted clone-related metrics and overall clonal architecture metrics between responder and non-responder samples. Positive values indicate higher values in the responder group, whereas negative values indicate higher values in the non-responder group. The first two metrics reflect the overall scale of highlighted clones and the maximum model score, whereas the last three metrics reflect the overall proportion of expanded clones, the maximum clone fraction, and clonal concentration within each sample. **(F)** Heatmap of mean expression differences between highlighted and other clones. See also Figure S5.

Consistent with previous studies linking the functional states of tumor-infiltrating CD8 T cells to response to PD-1 blockade^26^, the GEX-only model achieved relatively strong predictive performance. The TCR-only model performed substantially worse, indicating that TCR-derived information alone was insufficient for accurate cross-patient prediction. ImmuneLens outperformed both unimodal models across AUROC, AUPRC, balanced accuracy, and F1 score (**Figure 5b**). The fused model also achieved higher out-of-fold AUROC and AUPRC than the GEX-only model, with positive gains in paired patient-level bootstrap analyses (**Figure S5a,b**). We next examined the contribution of individual clones to each sample-level prediction.

Clone-level contributions followed a long-tailed distribution, with a small subset of top-ranked clones accounting for a disproportionate share of the prediction score (**Figure 5c**). We therefore defined the top 10% of clones within each sample as highlighted clones for subsequent biological characterization. Compared with other clones, highlighted clones had larger clone sizes and a higher fraction of TCR-expanded clones (**Figure 5d**). These differences remained evident across a range of highlighted-clone thresholds and were observed in most patients (**Figure S5c,d**). Metrics derived from highlighted clones showed stronger responder-associated differences than global clonal architecture metrics (**Figure 5e**). The predictive signal was therefore concentrated in a subset of response-associated clones rather than in the overall degree of clonal expansion.

Finally, highlighted clones showed higher expression of cytotoxicity- and activation/exhaustion-associated genes^26^ and lower expression of naive/memory genes than other clones (**Figure 5f**). These clones were characterized by clonal expansion together with cytotoxic and activation/exhaustion transcriptional programs, providing candidate clone-level features associated with immunotherapy response.

## DISCUSSION

T cell immune function depends not only on its current transcriptional state but is also influenced by prior antigen recognition and clonal selection processes. Single-cell transcriptomics can characterize T cell activation, differentiation, effector, and exhaustion states. TCRs and their clonal expansion patterns provide information on receptor specificity and clonal history. These two types of information are related but cannot substitute for each other. In this study, we incorporated GEX and TCR into a unified representation learning framework, which preserves cellular states and clonal backgrounds while further resolving T cell heterogeneity that is difficult to distinguish through unimodal analysis. Therefore, the value of ImmuneLens lies not only in improving downstream task performance, but also in the joint representation of transcriptional states and TCR clonal backgrounds.

Existing methods for the joint analysis of GEX and TCR address the relationship between the two modalities from different perspectives. CoNGA identifies cross-modal associations by comparing neighborhood structures in transcriptomic and TCR spaces^13^; Tessa focuses on combining TCR sequences with transcriptional states to characterize TCR functional networks^14^; deep generative models such as scNAT and mvTCR integrate paired GEX and TCR information into a joint latent representation^15,16^; MIST further constructs GEX, TCR, and joint representation spaces and emphasizes model interpretability^17^. Compared with these methods, ImmuneLens explicitly distinguishes GEX-private representations, TCR-private representations, and cross-modal shared representations, and establishes interactions between gene expression features and TCR sequence features through cross-attention. This design aims to reduce the dominance of a single modality over the fused representation and to provide a clearer structural basis for distinguishing shared information from modality-specific information.

In the antigen recognition and immunotherapy analyses, GEX and TCR provided complementary information that depended on the specific data context. The relative contributions of the two modalities differed across donors, while the fused representation showed better stability under class-imbalanced conditions. In the immunotherapy cohorts, joint analysis further distinguished T cell populations with similar transcriptional states but different associations with pathological response and different clonal backgrounds, and showed that sample-level predictive contributions were mainly concentrated in a subset of clones with both clonal expansion and cytotoxicity/activation-exhaustion programs. These results show that jointly analyzing T cell functional states and clonal backgrounds is more helpful for resolving antigen- and treatment-related immune heterogeneity than relying on either modality alone.

This study still has several limitations. First, ImmuneLens depends on paired single-cell GEX and TCR data, while the coverage of such data across different diseases, tissues, and treatment cohorts remains limited. The antigen recognition task is also constrained by the number of annotated samples, antigen diversity, and donor number. Therefore, the donor-specific modality contributions and cross-cohort generalizability observed in this study still need to be validated in larger and more diverse independent datasets. Second, the fused subclusters in LUAD and the immunotherapy-associated clones were mainly identified through retrospective computational analyses. Their stability, biological functions, and clinical significance still require further confirmation through independent cohorts, longitudinal sampling, and functional experiments. In addition, pathological response was used as the treatment outcome in this study and may not fully represent long-term survival benefit.

Overall, ImmuneLens provides a computational framework for the joint representation of T cell transcriptional states, TCR sequences, and clonal structures. By integrating the two data sources while preserving modality-specific information, this framework provides a new analytical perspective for characterizing T cell heterogeneity associated with antigen recognition and immunotherapy.

## RESOURCE AVAILABILITY

### Lead Contact

Further information and requests for resources should be directed to and will be fulfilled by the Lead Contact, Shuhui Song.

### Materials Availability

This study did not generate new unique reagents.

### Data and Code Availability

All datasets used in this study are publicly available. The colorectal cancer, non-small cell lung cancer, COVID-19, and basal cell carcinoma datasets were downloaded from NCBI GEO under accession numbers GSE236581, GSE243013, GSE158055, and GSE123813, respectively. The 10x Genomics dataset was obtained from the 10x Genomics website (https://www.10xgenomics.com/datasets).

All original code developed for this study is available at https://github.com/ddzzjj/ImmuneLens.

Any additional information required to reanalyze the data reported in this paper is available from the Lead Contact upon request.

## ACKNOWLEDGMENTS

We thank the Beijing Institute of Genomics, Chinese Academy of Sciences / China National Center for Bioinformation for providing computational resources. We also extend our gratitude to the researchers who generated the datasets and made them publicly available for this study. This work was supported by the National Natural Science Foundation of China (Grant No. 92374201).

## AUTHOR CONTRIBUTIONS

Z.D. and S.S. conceived the study, designed the figures, and wrote the manuscript. Z.D. developed the method, wrote the ImmuneLens code, and curated the experimental data. Y.W. contributed to experimental design and manuscript revision. C.L. revised the manuscript. G.L. contributed to conceptual design. Y.C. contributed to methodology design. X.B. and F.Y. provided input on experimental design. S.S. supervised the research and acquired funding. All authors read and approved the final manuscript.

## DECLARATION OF INTERESTS

The authors declare no competing interests.

## STAR METHODS

### Data preprocessing

For single-cell transcriptomic (scRNA-seq) data, standard preprocessing was performed using Scanpy^29^. Raw gene expression counts were normalized using normalize_total followed by log1p transformation, and the top 2,000 highly variable genes (HVGs) were selected using the Seurat v3 method with dataset as the batch key ^30^. Finally, the expression matrix was z-score scaled and clipped to the range [−10, 10]. For T cell receptor (TCR-seq) data, the CDR3β and CDR3α amino acid sequences of each cell were extracted. Only the 20 standard amino acid characters were retained, and the <start>, <end>, <pad>, and <mask> tokens were added. Sequences were truncated or padded to a fixed length of L = 20. CDR3β was used as the anchor chain, whereas the CDR3α chain was used as paired context, allowing TRA to be missing in some cells.

### Gene Encoder

To efficiently model high-dimensional and sparse gene expression data, we designed a hybrid gene encoder that combines a residual MLP with attention-based aggregation. The expression vector was first processed by LayerNorm and linear projection to obtain a global representation, *z*_global_. Meanwhile, the top-K = 512 genes were selected according to the absolute expression values, and sparse gene tokens were constructed by multiplying their expression values with learnable gene embeddings. These tokens were then aggregated by an attention aggregation module based on the Perceiver architecture ^31^ (n_latents = 64, n_heads = 8) to obtain a local key-gene representation, *z*_local_. The two representations were concatenated along the channel dimension, followed by linear fusion and further nonlinear transformation using a multilayer residual MLP with GEGLU activation, yielding the final GEX representation:

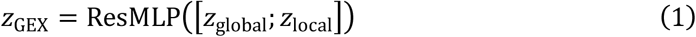

### TCR Encoder

A pretrained TCR language model, TCR-BERT, was used as the TCR sequence encoder. CDR3β and CDR3α sequences were separately input into independent TCR-BERT encoders to obtain token-level contextual embeddings. During training, most parameters were frozen, and only the last two Transformer Encoder layers of the CDR3β encoder were unfrozen for fine-tuning, while the CDR3α encoder remained frozen throughout training. This design preserved pretrained knowledge while preventing low-frequency TRA information from dominating training. The encoder output token-level embeddings and the corresponding valid-position masks, and sequence-level representations were obtained through mask-aware average pooling. In the antigen recognition prediction task, the protein language model ESM2 was also used as a TCR encoder, with parameter settings and training kept consistent with those of TCR-BERT for comparison.

### Construction of shared and private latent spaces

To simultaneously resolve modality-specific information and cross-modal covariation, ImmuneLens constructed GEX-private vectors, TCR-private vectors, and shared vectors. All three types of latent variables were generated through a VAE probabilistic bottleneck: *μ* and *log* σ^2^ were predicted separately, reparameterized sampling *z* = *μ* + *σ* ⊙ *ε* was used during training, and *μ* was used as the deterministic representation during inference. The GEX-private vector was derived from *z*_GEX_ output by the Gene Encoder and was used to preserve transcriptome modality-specific information. The TCR-private vector was generated from the cell-level *z*_TCR_, which was obtained by concatenating the pooled TRB representation with the TRA pooled representation weighted by α_*w*_ (default 0.3), followed by nonlinear mapping:

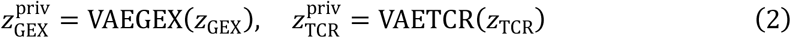

The KL weights of the two private bottlenecks in the total loss were additionally multiplied by 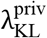 (default 2.0) relative to the shared bottleneck, to impose stronger constraints on modality-specific channels.

Shared features were used to model cross-modal associations between GEX and TCR. The model first concatenated TRB tokens with TRA tokens scaled byα_*w*_ along the sequence dimension to form a unified TCR token sequence T, and constructed the corresponding padding mask. This sequence was first passed through a layer of multi-head self-attention to enable intra-chain and inter-chain contextual exchange. During fusion, LN(*z*_GEX_) was used as a single query, while *LN*(*T*), attenuated by a TCR scale factor, was used as the key/value. Multi-head attention with the padding mask was then applied to obtain the contextual correction term Δ_TCR_, followed by LayerNorm normalization. To avoid excessive injection of TCR information, a scalar gate *g* was introduced at the fusion stage, which was predicted by a lightweight MLP and obtained through a sigmoid function. The final fused representation was passed through the shared VAE bottleneck to obtain *z*_shared_:

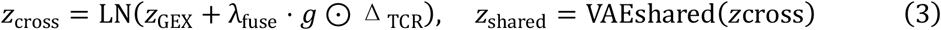

To reduce redundancy between the shared and private spaces, the model adopted an HSIC-based disentanglement constraint with an RBF kernel, where the kernel bandwidth was adaptively estimated from the mean pairwise distance within each batch. This constraint was applied between *z*_shared_ and 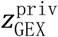, and between *z*_shared_ and 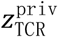, respectively:

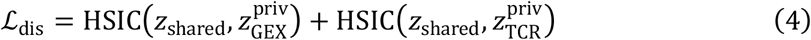

### Decoder design

GEX reconstruction adopted a design in which shared and private dual decoders were complementarily added in the output space. Both decoders were FiLM-conditioned residual MLPs^32^: the shared branch took *z*_shared_ as input, and the private branch took 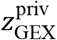 as input. In each FiLM residual block, the same latent vector was used to generate the scaling parameter γ through independent FiLM heads, which was bounded by tanh and multiplied by 0.5, and the shifting parameter β, which was multiplied by 0.5. The outputs of the two decoders were added element-wise along the gene dimension. The shared branch included a learnable output bias to learn the global gene mean, whereas the private branch disabled the bias to avoid redundancy:

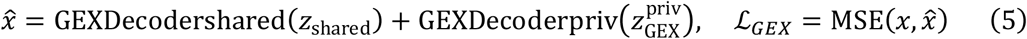

TCR sequence reconstruction similarly adopted a shared–private complementary design, but the outputs were added in the logit space. The model constructed two sets of autoregressive Transformer CDR3 decoders^33^ for TRB and TRA, respectively (pre-LN, n_layers = 2, n_heads = 4, d_ff = 512). The shared decoder used *z*_shared_ as a single-token memory, whereas the private decoder used 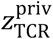 as a single-token memory. The two decoders used the same target token sequence, causal mask, and padding mask, and were trained with Teacher Forcing. The output logits were added and then passed through softmax to obtain token-wise prediction distributions. Token embeddings and output projections shared weights to reduce the number of parameters. TRB was used as the primary reconstruction target, whereas TRA was used as an auxiliary target with weight α (default 0.3), and the loss was computed only for cells with valid TRA:

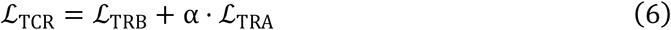

### Adversarial batch correction

During the reference atlas pretraining stage, we introduced gradient reversal-based domain-adversarial learning^34^. After the GEX representation, a spectral normalization-based residual MLP batch discriminator *D* was constructed to predict the dataset origin of each cell. The encoder and discriminator were connected through a gradient reversal layer (GRL), enabling the discriminator to minimize the batch classification error while the encoder maximized this error through reversed gradients, thereby learning batch-insensitive latent representations:

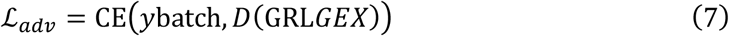

The adversarial strength λ_*adv*_ was smoothly increased from 0 to an upper bound λ_*max*_ (default 0.3) during training according to a sigmoid schedule.

### Integration performance evaluation

We evaluated the cross-dataset integration performance of different methods using the Overall integration score, Adjusted Rand Index (ARI), Normalized Mutual Information (NMI), and batch average silhouette width (ASW batch). ARI and NMI were used to measure the agreement between the clustering results in the integrated space and the annotated cell-type labels, with higher values indicating better preservation of cell-type structure. ASW batch was used to assess the mixing of cells from different datasets within the same cell type, with higher values indicating more effective batch-effect correction.

The Overall integration score jointly considers biological structure preservation and batch-effect correction. The biological conservation score was calculated as follows:

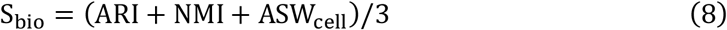

The batch-correction score was calculated as follows:

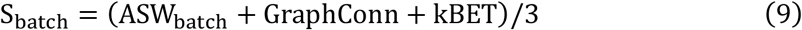

The Overall integration score was calculated as follows:

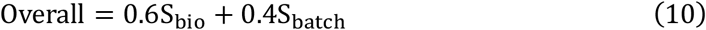

A higher Overall integration score indicates that the method achieves a better balance between preserving cell-type-related biological structure and reducing batch effects across datasets.

### Reference atlas construction and query mapping

External query data were processed using the same preprocessing workflow as the reference atlas. normalize_total followed by log1p transformation was applied, and genes were aligned to the HVG list saved during the reference stage, with missing genes filled with zeros. The query data were then mapped to the reference latent space using the pretrained Gene Encoder with frozen parameters. To preserve the global topology of the reference atlas, the latent representations of query cells were reduced using the PCA model saved during the reference stage. Then, k nearest neighbors were searched in the reference space using cosine distance, and the distances were converted into exponential weights. The two-dimensional UMAP coordinates of query cells were obtained as the weighted average of the UMAP coordinates of their neighboring reference cells, without rerunning UMAP. Annotation transfer was based on the cosine similarity between the latent centroid of each reference cell type and the query representation, with the Top-1 prototype selected as the predicted label. Prediction confidence was defined as this Top-1 cosine similarity, and projection uncertainty was defined as the average distance between the query cell and its k nearest reference cells. These two metrics were jointly used to identify low-confidence or previously uncharacterized cell states.

### Overall training objective

The overall training objective of the model consisted of GEX reconstruction, TCR reconstruction, shared–private disentanglement, KL regularization of the probabilistic latent space, and the adversarial loss, which was enabled only during reference atlas pretraining:

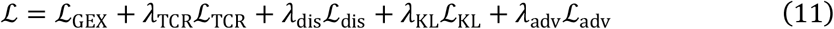

Here, ℒ_KL_ denotes the sum of the KL terms from the three branches, *z*_shared_, 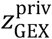, and 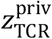, with the two private branches additionally multiplied by 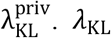 was applied with a linear warm-up strategy to prevent overly strong latent regularization from suppressing reconstruction during the early stage of training. In the antigen recognition prediction task, the pretrained encoder was frozen, and the cell-level fused embedding was obtained by independently applying L2 normalization to *z*_shared_, 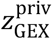, and 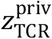, concatenating them, and then applying overall L2 normalization. A lightweight paired classification head was trained for each donor, in which the cell embedding and antigen embedding were concatenated and passed through an MLP to output logits. A BCE loss weighted by pos_weight was used, and early stopping was performed based on validation AUPRC.

### Clone-aware MIL framework for clinical response prediction

In NSCLC neoadjuvant ICB pathological response prediction, we combined *z*_shared_, 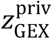, and 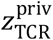 with multiple instance learning (MIL). Each bag consisted of a set of paired GEX+TCR single cells from the same tumor sample (sample-level) or the same patient (patient-level), and the bag label corresponded to the pathological response. The GEX-only and TCR-only baselines used the same VAE bottleneck and MIL head, with only the encoder replaced. By default, the fused model used 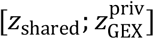 as the clonal representation input. Evaluation was performed using an outer StratifiedGroupKFold cross-validation grouped by PatientID. GEX preprocessing steps, including HVG selection, log1p transformation, and scaling, were fitted only on the training bags within each fold and then applied to the validation/test bags to avoid information leakage.

## Notes

### Competing Interest Statement

The authors have declared no competing interest.

