## Supplemental Information for "ImmuneLens: linking transcriptional states and TCR clonotypes through disentangled multimodal learning"

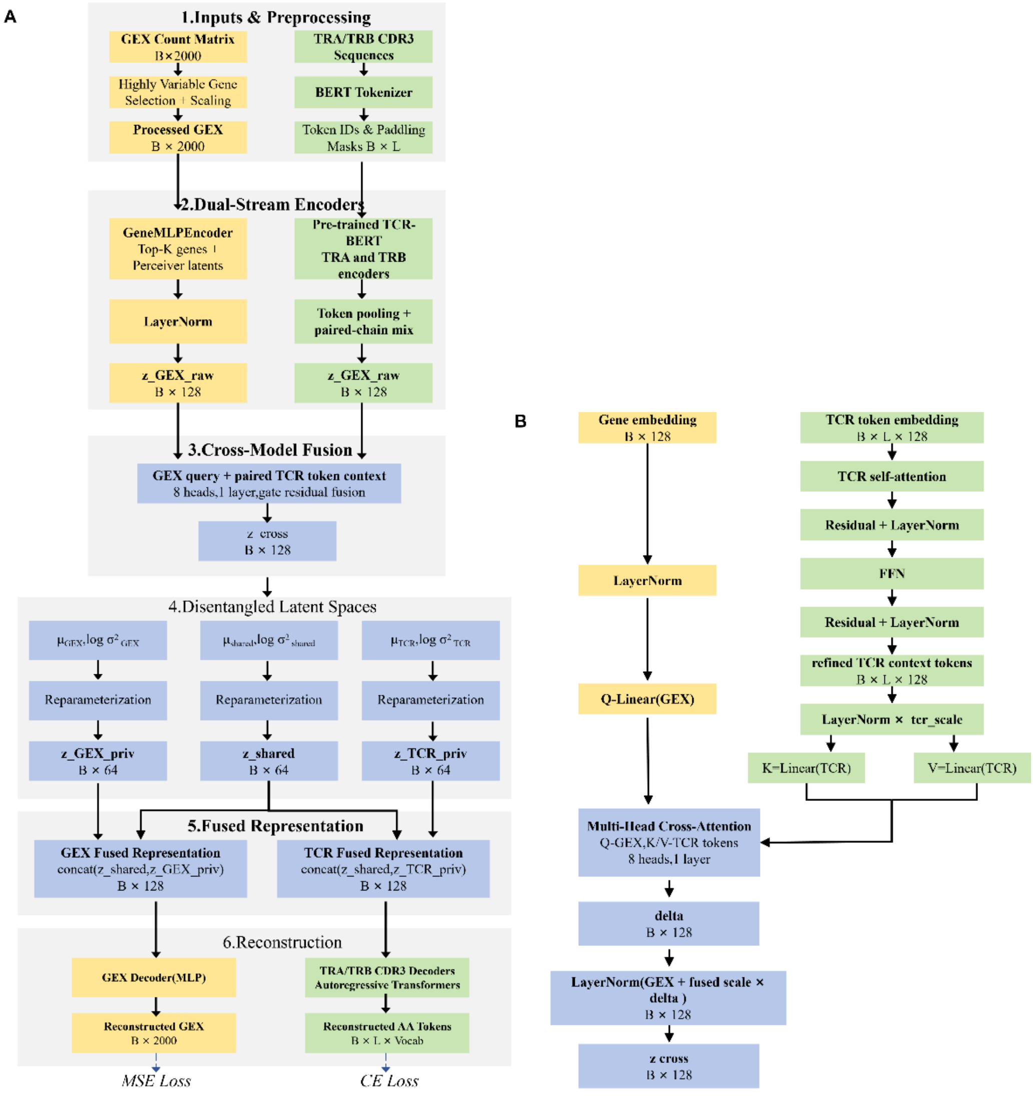


**Figure S1. Detailed architecture of the ImmuneLens model and cross-modal attention module, Related to Figure 1**

**A.** Detailed network architecture of ImmuneLens. Gene expression count matrices are preprocessed by highly variable gene selection, normalization, and scaling, and are subsequently encoded by a hybrid gene encoder combining global gene expression features with attention-based aggregation of highly expressed genes. Paired TRA and TRB CDR3 sequences are tokenized and processed by pretrained TCR-BERT encoders to obtain token-level and sequence-level TCR representations. The GEX representation attends to the paired TCR token context through multi-head cross-attention, producing a cross-modal representation. GEX-private, TCR-private, and cross-modal shared latent variables are generated through separate variational bottlenecks, with HSIC-based constraints used to reduce redundancy between shared and private spaces. The shared and private representations jointly support GEX and TCR sequence reconstruction.

**B.** Detailed structure of the cross-modal attention module. Paired TRA/TRB token embeddings are first refined through TCR self-attention and a feed-forward network. The normalized GEX representation is projected as the query, whereas the refined TCR tokens are projected as keys and values. Multi-head cross-attention produces a TCR-derived contextual correction term, which is incorporated into the GEX representation through gated residual fusion and LayerNorm to generate the cross-modal representation.

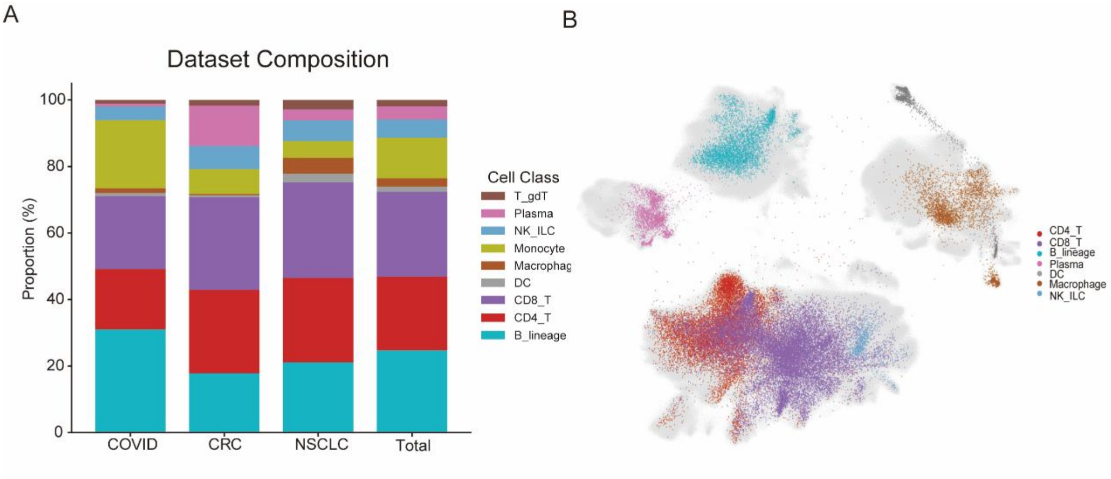


**Figure S2. Cell type composition of the multi-cohort reference datasets and the external BCC query dataset, Related to Figure 2**

**A.** Proportions of major immune cell types in the COVID-19, colorectal cancer (CRC), and non-small cell lung cancer (NSCLC) datasets used to construct the multi-cohort immune reference atlas. “Total” indicates the overall cell type composition after combining the three reference datasets.

**B.** UMAP visualization of the external basal cell carcinoma (BCC) query dataset, colored by major immune cell type.

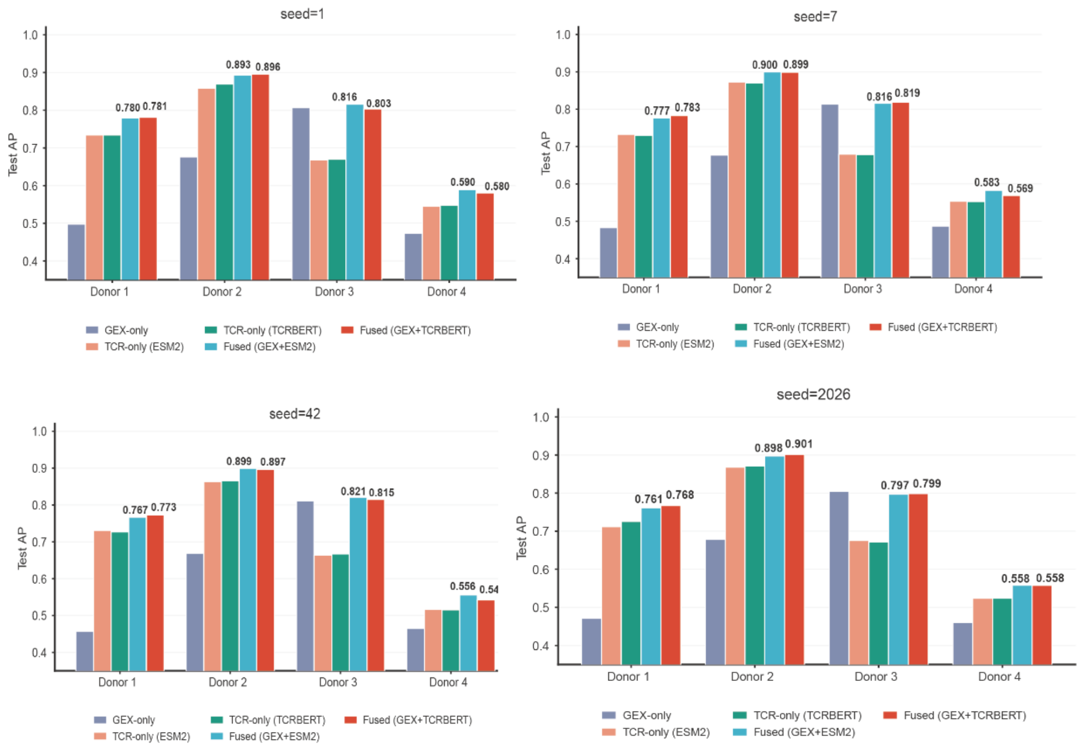


**Figure. S3. Robustness of antigen recognition performance across different random seeds, Related to Figure 3**

Test average precision (AP) of the GEX-only, TCR-only, and fused models across four donors under four random seeds (1, 7, 42, and 2026). TCR-BERT and ESM2 were used as alternative TCR sequence encoders. Across different random initializations, the fused models generally maintained competitive or superior performance relative to the corresponding unimodal baselines, indicating that the antigen recognition results were robust to random seed variation.

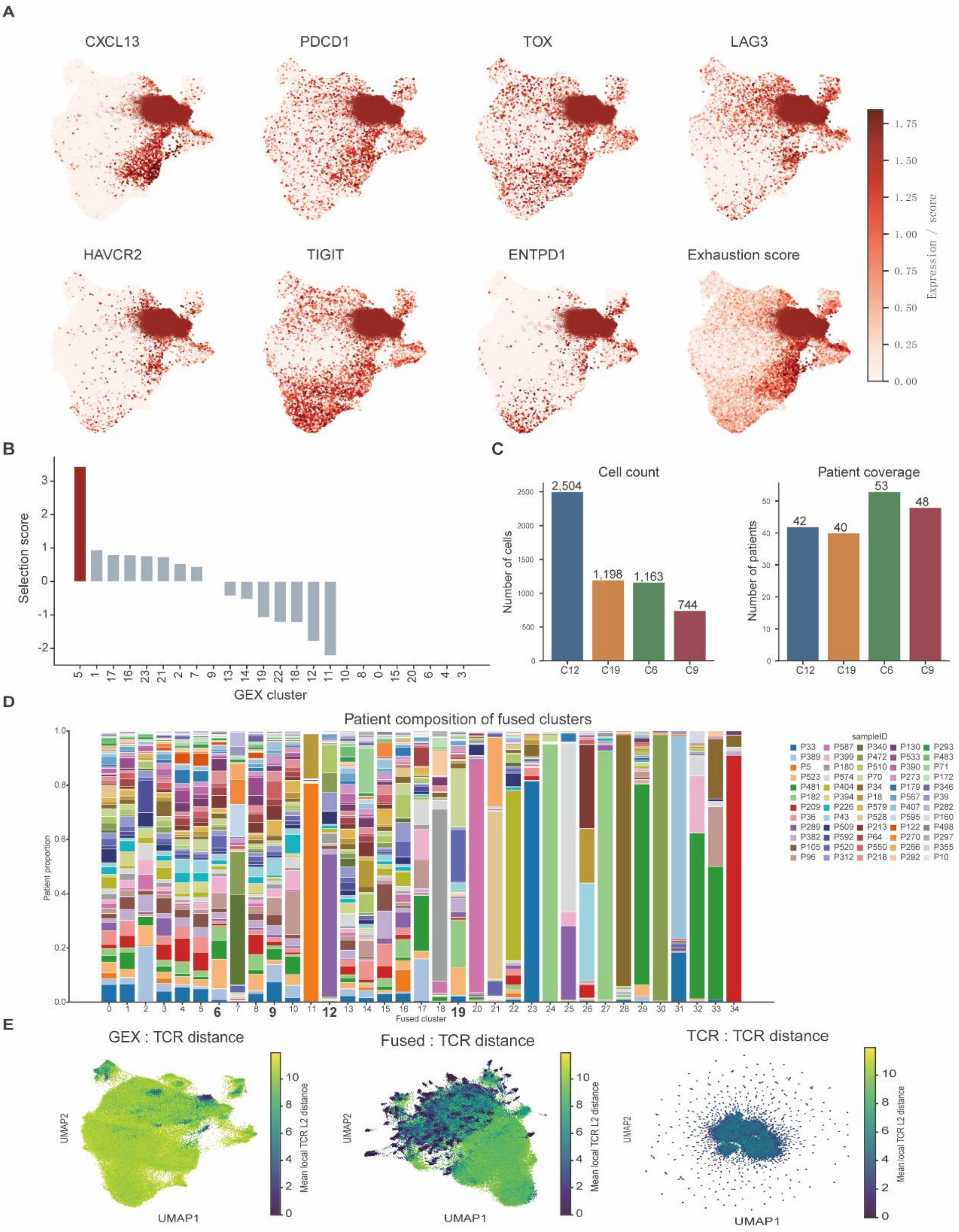

A
CXCL13
PDCD1
TOX
LAG3
HAVCR2
TIGIT
ENTPD1
Exhaustion score
Expression / score
0.00
0.25
0.50
0.75
1.00
1.25
1.50
1.75
B
Selection score
GEX cluster
5 1 17 16 23 21 2 7 9 13 14 19 22 18 12 11 10 8 0 15 20 6 4 3
C
Cell count
Number of cells
2,504
1,198
1,163
744
C12 C19 C6 C9
Patient coverage
Number of patients
42
40
53
48
C12 C19 C6 C9
D
Patient composition of fused clusters
Patient proportion
Fused cluster
sampleID
P33 P587 P340 P130 P293
P389 P399 P472 P533 P483
P5 P180 P510 P390 P71
P523 P574 P70 P273 P172
P481 P404 P34 P179 P346
P182 P394 P18 P567 P39
P209 P226 P579 P407 P282
P36 P43 P528 P595 P160
P289 P509 P213 P122 P498
P382 P592 P64 P270 P297
P105 P520 P550 P266 P355
P96 P312 P218 P292 P10
E
GEX : TCR distance
Fused : TCR distance
TCR : TCR distance
UMAP1
UMAP2
Mean local TCR L2 distance

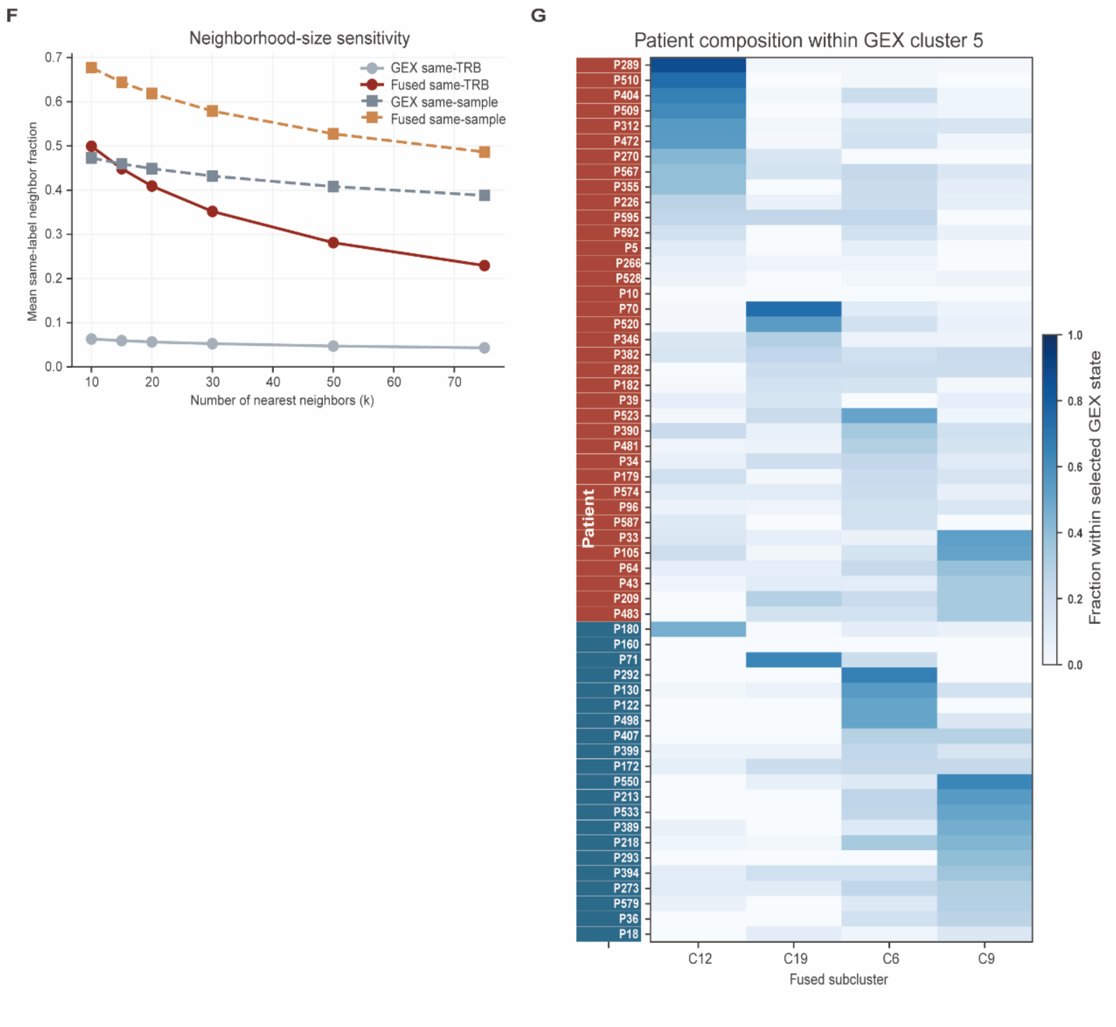


**Figure S4. Selection, patient composition, and TCR-associated organization of fused subclusters in the LUAD cohort, Related to Figure 4**

**A.** UMAP visualization of tumor-reactive/exhaustion-associated genes, including CXCL13, PDCD1, TOX, LAG3, HAVCR2, TIGIT, and ENTPD1, together with the composite tumor-reactive/exhaustion score in the GEX representation space.

**B.** Ranking of GEX clusters according to the composite selection score. GEX cluster 5 achieved the highest score and was selected as the tumor-reactive/exhaustion-like state for subsequent fused-representation analysis.

**C.** Cell counts and patient coverage of the four major fused subclusters within GEX cluster 5, including C12, C19, C6, and C9.

**D.** Patient composition of individual fused clusters within GEX cluster 5. Each stacked bar represents one fused cluster, and colors indicate the relative contributions of individual patients.

**E.** Distribution of local TCR distances in the GEX, fused, and TCR representation spaces. Cells are colored by their mean local TCR distance, with lower values indicating greater local similarity in TCR-derived features.

**F.** Neighborhood-size sensitivity analysis of local TRB clonotype and patient consistency in the

GEX and fused representation spaces. The mean fraction of nearest neighbors sharing the same TRB clonotype or originating from the same patient was evaluated across different neighborhood sizes.

**G.** Patient-level abundance of the four major fused subclusters within GEX cluster 5. Rows represent patients and columns represent fused subclusters. Values indicate the fraction of cells assigned to each fused subcluster within the selected GEX state. The row-side annotation indicates pathological response group.

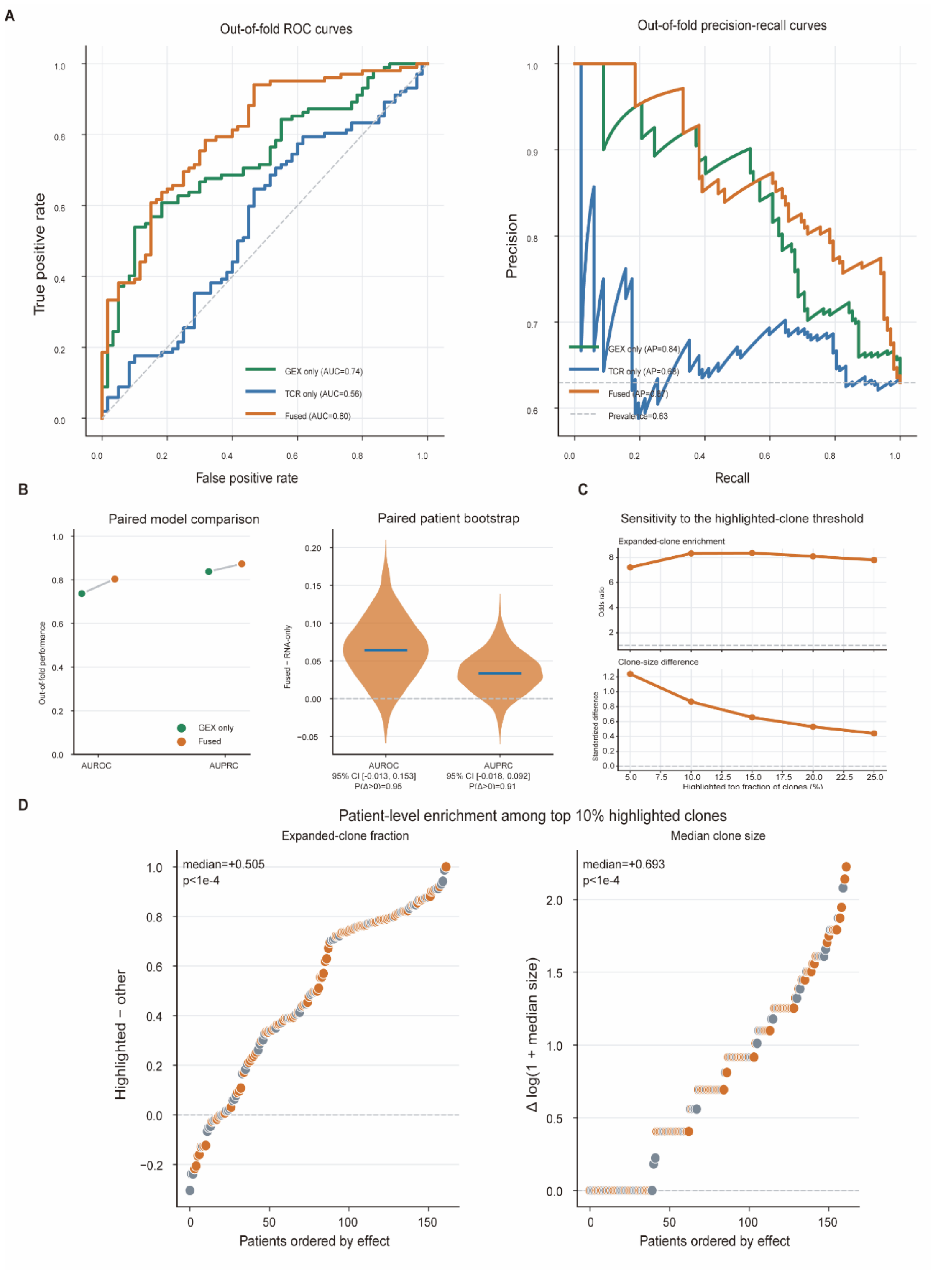


**Figure S5. Performance robustness and clonal enrichment analyses for immunotherapy response prediction, Related to Figure 5**

**A.** Out-of-fold ROC and precision–recall curves for the GEX-only, TCR-only, and fused models.

**B.** Paired comparison of out-of-fold AUROC and AUPRC between the GEX-only and fused models, together with paired patient-level bootstrap distributions of the performance gain of the fused model.

**C.** Sensitivity analysis of the highlighted-clone threshold. Expanded-clone enrichment and clone-size differences between highlighted and other clones were evaluated across different top-fraction thresholds.

**D.** Patient-level differences in expanded-clone fraction and median clone size between the top 10% highlighted clones and other clones. Patients were ordered according to the magnitude of the corresponding effect.
